# PPARG governs adipogenic differentiation and cell state plasticity in well-differentiated and dedifferentiated liposarcoma

**DOI:** 10.1101/2025.07.16.665211

**Authors:** Blake R. Wilde, Kyle D. Klingbeil, Francesca Day, Chris Dann, Chris Frias, Manando Nakasaki, Sarah M. Dry, Fritz C Eilber, Joseph G. Crompton, David B. Shackelford, Brian E. Kadera, Heather R. Christofk

## Abstract

Well-differentiated and dedifferentiated liposarcoma (WD/DD LPS) represent a pathological continuum, often coexisting within the same tumor. While the dedifferentiated component is clinically aggressive, marked by rapid growth and metastatic potential, the evolutionary relationship between WD and DD LPS remains unknown. To investigate this, we performed single-nucleus RNA sequencing on matched WD and DD tumor regions. Both compartments shared a predominant population of undifferentiated mesenchymal cells, but only WD regions contained cells expressing adipocytic differentiation markers and PPARG target genes. Given the central role of PPARG in coordinating lipid metabolism and mitochondrial biogenesis during adipogenesis, these findings suggest that loss of this program may underlie the poorly differentiated, proliferative phenotype of DD LPS. Functional studies confirmed that PPARG activation in DD LPS cells induces lipid accumulation, reduces proliferation, and impairs tumor growth *in vivo*. These support a model in which impaired adipogenic differentiation underlies DD LPS pathology and identify PPARG as a potential therapeutic target to promote differentiation and suppress tumor progression.

**Teaser:** PPARG reprograms DD liposarcoma toward adipogenesis, reducing proliferation and tumor growth

## Introduction

Liposarcoma (LPS) is the most common soft tissue sarcoma subtype in adults, and the incidence is rising(*1, 2*). Understqanding of its biology lags behind other malignancies and treatment options remain limited(*3–5*). Surgical resection is the primary treatment, but recurrence occurs in more than 50% of cases and there are currently no curative options for recurrent disease (*6–8*). These challenges underscore the urgent need for novel therapeutic strategies.

The World Health Organization (WHO) classifies LPS into five major subtypes: well-differentiated (WD), dedifferentiated (DD), myxoid, pleomorphic, and myxoid-pleomorphic. Among these, WD and DD LPS are the most prevalent and are increasingly recognized as a spectrum of the same disease—collectively termed WD/DD LPS—due to shared defining genomic feature: amplification of the 12q13-15 locus, including *MDM2*(*9, 10*).

Clinically, WD/DD LPS manifests in various forms: as a homogeneous WD or DD LPS tumor, or as a tumor with distinct components of <u>both</u> WD and DD LPS (*11, 12*). Although WD LPS is generally indolent, some cases progress rapidly to DD LPS(*13*). Conversely, DD LPS— while generally associated with a high metastatic potential and a six-fold increased risk of death—can occasionally follow a more protracted course(*14–18*). The observation that tumors can recur with a histology different from the original subtype, including instances of DD LPS recurring as WD, suggests that tumor differentiation exists along a dynamic continuum rather than as a fixed binary state(*12*). These clinical scenarios suggest a dynamic, bidirectional differentiated-dedifferentiated state, but the underlying molecular mechanisms remain poorly understood. This clinical heterogeneity also indicates that histologic classification alone is insufficient for the prognostication that drives treatment decisions. Molecular markers such as MDM2 amplification level and IGF2BP3 expression have shown improved prognostic value, emphasizing the need for a deeper understanding of tumor evolution in WD/DD LPS (*19, 20*).

Histologically, WD LPS is composed of mature adipocytes and lipoblasts with nuclear atypia, while DD LPS consists of undifferentiated spindle cells with high mitotic rate and reduced expression of adipogenic markers (*7, 13, 21*). Both subtypes are thought to originate from mesenchymal precursors such as adipocyte progenitor cells or preadipocytes. Genomic analyses have revealed similar somatic mutation profiles between WD and DD LPS, yet DD tumors exhibit greater genomic complexity, including additional copy number alterations beyond 12q13–15— such as amplifications at 5p and 14q and deletions at 11q23–24, 19q13, 3q29, 9p22–24, or 17q21—suggesting secondary genomic events may drive dedifferentiation(*22–26*).

Viewing WD/DD LPS as a dynamic spectrum provides a framework to improve diagnosis, prognostication, and therapeutic development. Although bulk genetic profiling has yielded valuable insights, it has yet to elucidate the molecular mechanisms that govern differentiation state and tumor progression. Single-cell and spatial omics approaches offer a powerful means to dissect intra-tumoral heterogeneity and trace tumor evolution. A recent study by Gruel *et al.* used single-cell transcriptomics to demonstrate that both WD and DD components originate from a shared adipocyte stem cell and that differentiation in DD LPS is suppressed by TGF-β signaling(*27*).

Building on this work, we performed single-nucleus RNA sequencing (snRNA-seq) on paired WD and DD tumor components from patients with WD/DD LPS. Our data revealed significant inter-patient heterogeneity, but a shared core architecture: both WD and DD compartments were predominantly composed of undifferentiated mesenchymal cells, with only WD regions containing cells expressing markers of preadipocytes and mature adipocytes. Pseudotime trajectory analysis revealed a clear transcriptional path toward adipogenic differentiation in WD tumors, associated with activation of PPARG target genes.

These findings led us to test whether PPARG activation could reprogram DD LPS cells toward a more differentiated, less aggressive state. We found that the splice variant PPARG2 promoted lipid accumulation, reduced proliferation in vitro, and suppressed tumor growth in vivo. Moreover, pharmacologic activation of PPARG with rosiglitazone impaired the growth of DD LPS xenografts. Together, these results suggest that loss of PPARG-driven differentiation underlies dedifferentiation in LPS and that restoring this program may represent a promising therapeutic strategy.

## Results

### snRNA-seq identifies molecular heterogeneity in WD/DD LPS

The molecular basis of WD/DD LPS transition remains poorly understood, despite their frequent co-occurrence within the same tumor. This spatial juxtaposition provides a unique opportunity to investigate dedifferentiation in situ—offering direct insight into tumor evolution, plasticity, and progression. We hypothesized that distinct transcriptional programs and differentiation states underlie the histological differences between WD and DD components, and that single-nucleus RNA sequencing (snRNA-seq) could resolve these programs at high resolution. This approach is uniquely suited for LPS because it enables detection of mature adipocytes, which are typically lost in single-cell RNA-seq using droplet-based platforms(*28*). This is particularly important in LPS, where mature adipocytes and lipid-laden lipoblasts define WD histology and may represent a differentiation endpoint or lineage anchor. Capturing these populations allows for more accurate reconstruction of adipogenesis and identification of dedifferentiation events.

To identify tumors with clear histologic demarcation between WD and DD components, we screened primary, untreated WD/DD LPS cases based on CT imaging and confirmed subtype identity using histology and DNA-FISH (**Fig. 1A–C**). Three tumors were selected for analysis (**Table 1**). The tumors analyzed varied widely in size (8–46 cm) and clinical behavior. This heterogeneity reflects the clinical challenges posed by WD/DD LPS and underscores the need for a deeper molecular understanding. WD components showed abundant mature adipocytes and lipoblasts, while DD regions were composed of densely cellular, non-lipogenic spindle cells (**Fig. 1B**, **Fig. S1A–B**). *MDM2* amplification was confirmed in both regions, validating their malignant origin and enabling distinction between tumor and non-tumor nuclei in downstream analyses.

**Fig. 1.**
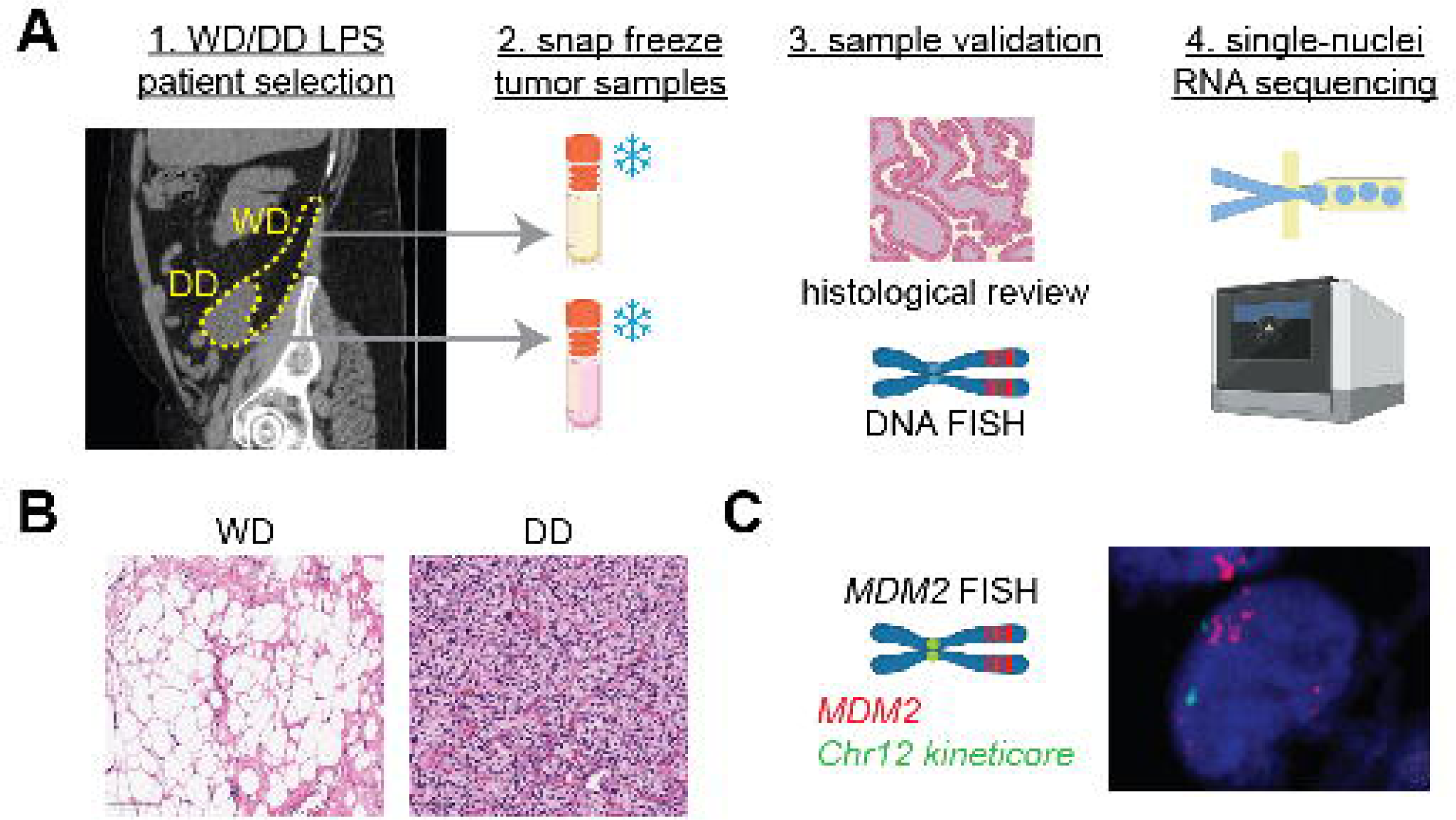
Study overview and histologic characterization of WD/DD liposarcoma for single-nucleus RNA sequencing. (**A**) Schematic of experimental workflow. Computed tomography (CT) scan from a patient with untreated retroperitoneal WD/DD LPS (sagittal view), demonstrating differential contrast enhancement consistent with spatially distinct WD and DD tumor regions. Tumor samples were collected at the time of surgical resection. Both WD and DD components were independently macrodissected from each tumor, analyzed for histological and molecular features, and subjected separately to snRNA-seq. (**B**) H&E staining of matched WD and DD components from the same tumor. The WD region is characterized by abundant lipoblasts and adipocytic differentiation, while the DD component shows increased cellularity and nuclear atypia. (**C**) DNA-FISH for MDM2 reveals gene amplification (red), with chromosome 12 centromere probe (green) as control.

**Table 1.**
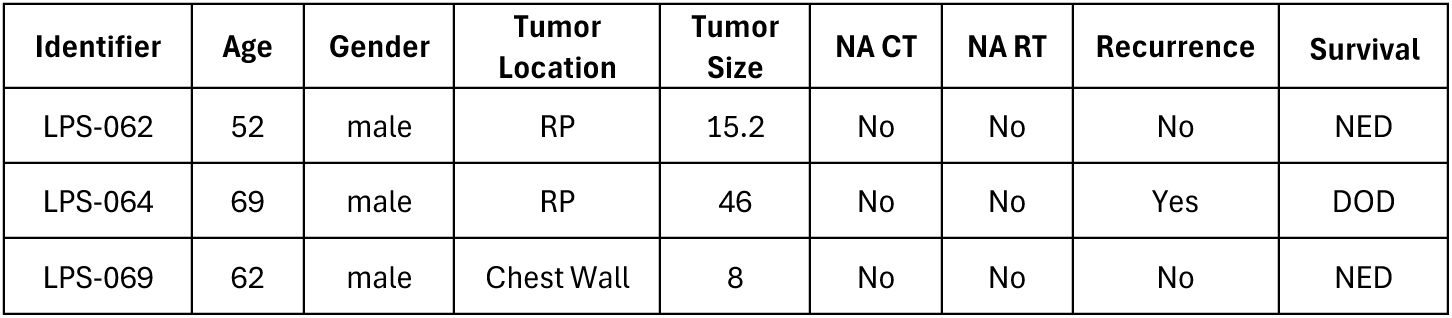
Clinicopathologic variables of WD/DD LPS tumor cohort. Tumor size is the greatest dimension in centimeters. CT, chemotherapy; DOD, dead of disease; NA, neoadjuvant; NED, no evidence of disease; RP, retroperitoneum; RT, radiation therapy. Median follow-up is 2.2 years.

Following tissue dissociation and nuclei extraction, cDNA libraries were constructed and sequenced. After read alignment and quality filtering using Cell Ranger and CellBender, we integrated datasets by patient and subtype using Seurat(*29–31*). Dimensionality reduction and clustering revealed 22 distinct transcriptional populations (**Fig. 2A**). Of these, 10 clusters expressed canonical LPS oncogenes (*MDM2*, *CDK4*, *HMGA2*) (**Fig. 2B**, **Fig. S2**), suggesting a tumor cell identity. To validate this classification, we performed copy number inference, which revealed characteristic gains on chromosome 12q13–15—encompassing the *MDM2* locus—specifically in these clusters (**Fig. 2C**). Inferred CNAs provided orthogonal confirmation of tumor identity and helped distinguish malignant from stromal and immune populations.

**Fig. 2.**
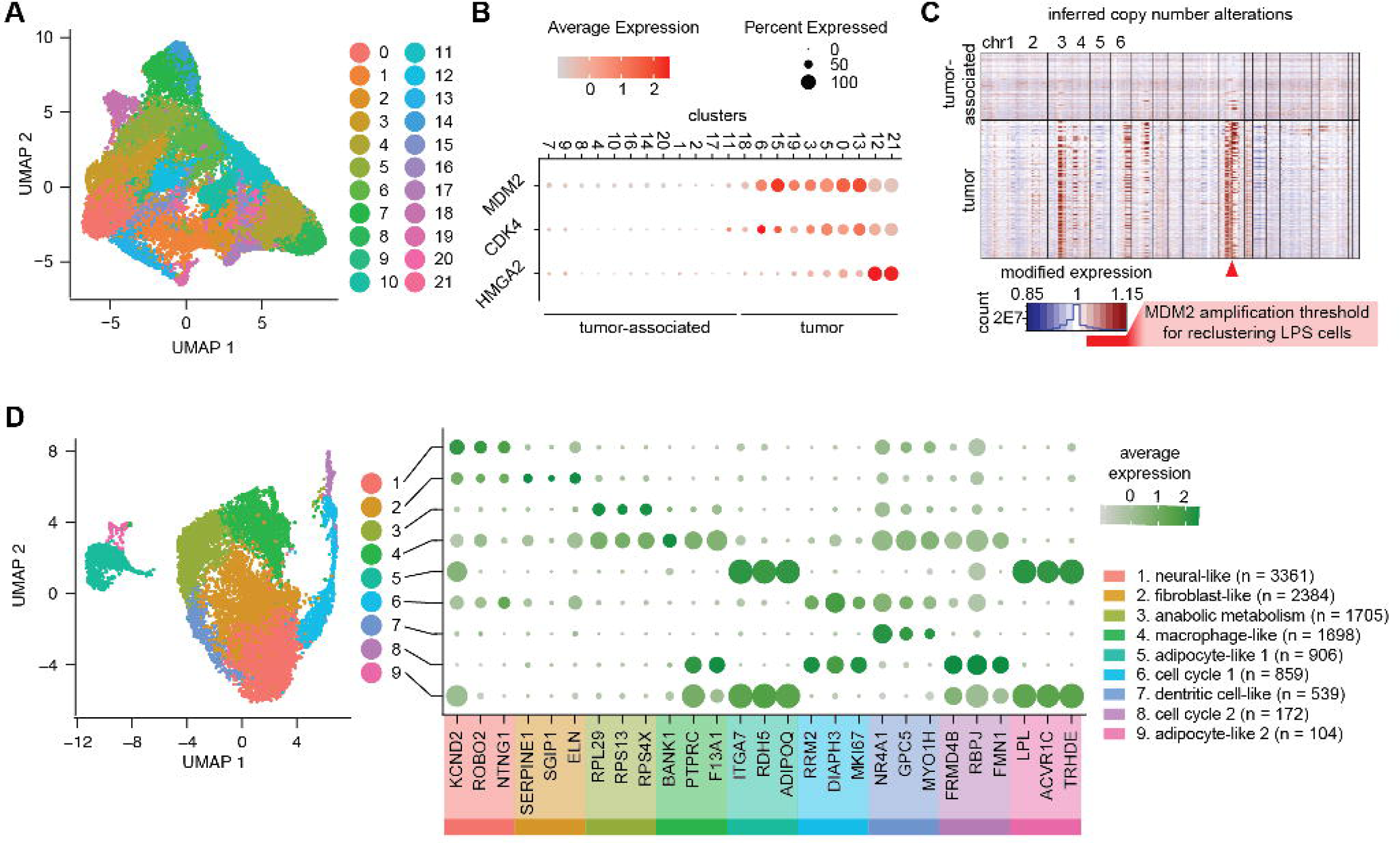
Single-nucleus transcriptomic profiling identifies malignant clusters and reveals transcriptional heterogeneity in WD/DD LPS. **(A)** UMAP embedding of all nuclei from WD and DD components across three patient tumors reveals 21 transcriptionally distinct clusters. (**B**) Expression of known LPS-associated genes was used to annotate and predict clusters enriched for malignant tumor cells. (**C**) Inferred copy number alterations (CNAs) show characteristic gains on chromosome 12q—encompassing the MDM2 locus—specifically in the predicted tumor cell clusters. (**D**) Re-clustering of tumor-enriched nuclei reveals nine malignant subclusters, each defined by distinct gene expression signatures and predicted functional states.

To focus our analysis on tumor-intrinsic heterogeneity, we re-clustered *MDM2*-amplified nuclei and identified nine transcriptionally distinct tumor cell states (**Fig. 2D**) (*32*). These included proliferative clusters, metabolically active populations, and cells with mesenchymal, immune-like, or adipogenic features (**Fig. S3A-D**). These data reveal extensive intra-tumoral diversity and support the existence of discrete tumor cell states that may influence growth, recurrence, and therapeutic response.

### WD tumors uniquely harbor differentiated adipocyte-like cells

We next asked whether specific tumor cell populations were preferentially enriched in either the WD or DD regions. Among the nine tumor-intrinsic clusters, two (clusters 5 and 9) stood out for their high expression of adipocyte lineage markers including *PPARG*, *FABP4*, and *ADIPOQ*, consistent with a differentiated adipogenic phenotype (**Fig. 3A-B**). Notably, these cells lacked expression of adipocyte progenitor markers such as *COL6A1*, *COL6A2*, and *FKBP10*, reinforcing the idea that they are committed to adipocyte fate rather than representing an intermediate precursor state (**Fig. 3C**). These adipocyte-like clusters accounted for approximately 9% of all tumor cells and were detected exclusively within the WD component, suggesting they may represent a terminally differentiated cell population (**Fig. 3D**, **Fig. S3A**). We refer to these as “WD-specific clusters.”

**Figure 3.**
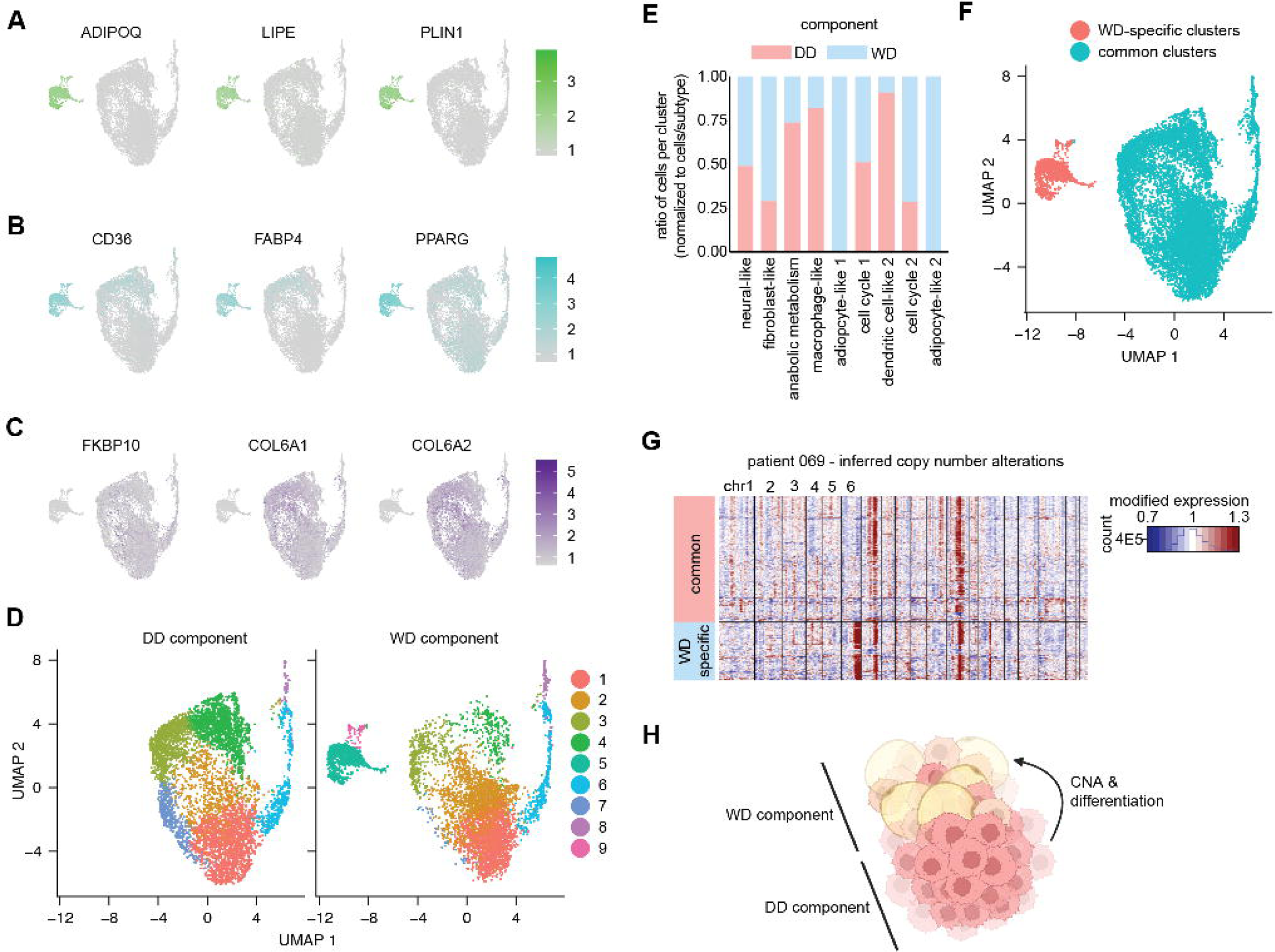
Adipocyte-like tumor cells are restricted to the WD component. UMAPs show expression of (**A**) mature adipocyte markers (ADIPOQ, LIPE, PLIN1), (**B**) preadipocyte markers (CD36, FABP4, PPARG), and (**C**) adipocyte progenitor markers (COL6A1, COL6A2, FKBP10), highlighting a distinct population of differentiated cells exclusive to the WD component. (**D**) UMAPs split by WD and DD components confirm the spatial restriction of this population. (**E**) Quantification of cluster composition by subtype shows that adipocyte-like clusters are present only in WD tumors. (**F**) UMAP colored by shared versus WD-specific cluster identity. (**G**) Inferred copy number profiles from patient 069 demonstrate shared CNAs (12q, 7q) across both components and additional gains (6q, 14q) restricted to the WD-specific cluster. (**H**) Model summarizing the differentiation trajectory from shared progenitor states to the WD-specific adipocyte-like cell population.

To determine whether other tumor cell populations were similarly restricted to one component or shared between both, we examined the distribution of all clusters across matched WD and DD samples. Most clusters were present in both components, including proliferative, mesenchymal-like, and metabolically active populations, indicating that despite histological divergence, the majority of tumor cells share a conserved transcriptional architecture (**Fig. 3E**). We refer to these as common clusters, as they exhibited similar abundance and transcriptional profiles in WD and DD regions (**Fig. 3F**). The presence of shared transcriptional states across spatially distinct histologies suggests that WD and DD tumors retain a core transcriptional architecture, and that differentiation states are not driven by global reprogramming but instead by selective enrichment or loss of specific states. The key distinction between components was the exclusive presence of the adipocyte-like, WD-specific clusters—a feature entirely absent from the DD regions.

To investigate the lineage relationship between tumor cell populations, we performed copy number inference on WD and DD tumor cells. In one representative tumor (patient #069), both WD and DD components shared canonical gains on 12q and 7q, consistent with a common clonal origin. However, only the WD-specific clusters harbored additional CNAs—including gains on 6q and 14q—suggesting these cells diverged from the shared progenitor through a distinct genetic trajectory (**Fig. 3G**). This supports a model in which both WD-specific and common clusters originate from a shared population of undifferentiated tumor cells, with differentiation governed by acquisition of CNAs and/or activation of lineage-specific transcriptional programs.

It is unclear whether WD and DD represent distinct, parallel tumor lineages or reflect a continuum in which one subtype gives rise to the other. Shared transcriptional architecture and copy number alterations support a common clonal origin. However, the presence of additional CNAs unique to the WD component—absent from DD—suggests the WD compartment may arise via differentiation from a less differentiated DD-like progenitor (**Fig. 3H**). In this model, WD tumor cells acquire new genetic features or engage specific transcriptional programs that enable adipogenic differentiation and give rise to the histologically well-differentiated phenotype. The complete absence of adipocyte-like cells in DD regions further supports the idea that differentiation, rather than dedifferentiation, explains the emergence of WD histology in some cases. This concept is supported by recent studies that show that both WD and DD components arise from a shared adipocyte stem cell–like progenitor(*27*). Understanding the mechanisms that constrain or permit this differentiation could provide new opportunities to manipulate tumor cell state with novel systemic therapies.

### Pseudotime analysis implicates PPARG in adipogenic differentiation

We next sought to identify the regulatory programs that govern the transition from undifferentiated tumor cells to adipocyte-like states. Given the restricted presence of adipogenic clusters in WD tumors, we hypothesized that WD/DD subtype divergence reflects altered transcriptional trajectories rather than the presence of entirely distinct lineages. To test this, we performed pseudotime trajectory analysis using Monocle 3, anchoring the trajectory at the WD-specific adipocyte-like clusters as a terminal node(*33*).

Trajectory reconstruction revealed a complex network of potential paths originating within the common clusters, which comprised transcriptionally heterogeneous populations with the capacity to adopt multiple fates. Within this network, we identified a continuous trajectory linking the common clusters to the WD-specific adipocyte-like clusters, suggesting a directed progression toward adipogenic differentiation (**Fig. 4A**). Along this path, we observed a gradual decrease in *MDM2* expression and a corresponding increase in *HMGA2*, two markers historically associated with DD and WD histology, respectively (**Fig. 4B**). These findings support a model in which the common clusters serve as a transitional hub, giving rise to either differentiated adipocyte-like cells—represented by the WD-specific clusters—or to alternative, non-lipogenic tumor states.

**Figure 4.**
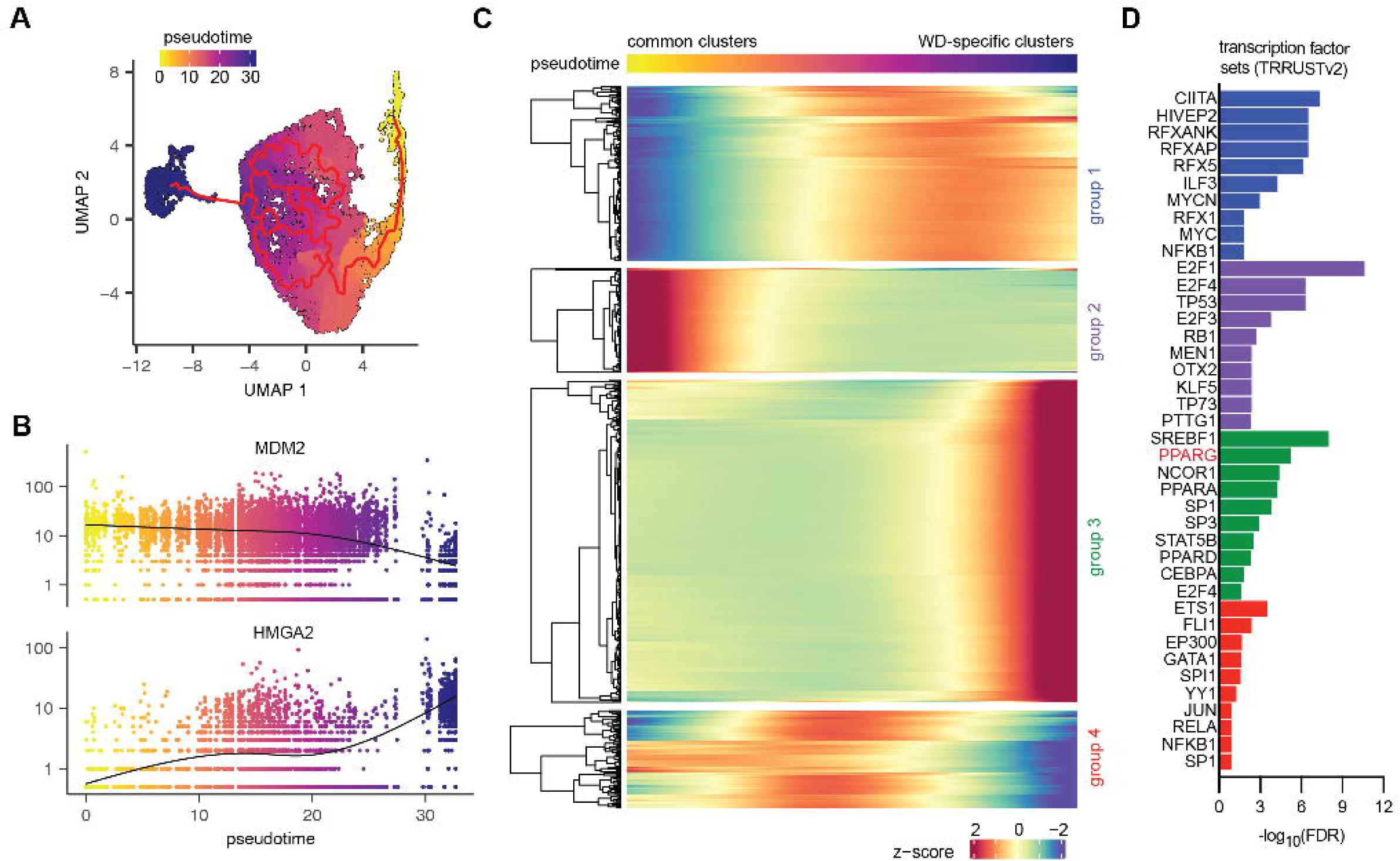
Pseudotime analysis reveals transcriptional trajectories toward adipogenic differentiation. (**A**) Monocle 3 trajectory analysis reconstructs a continuous path from common tumor clusters to the WD-specific adipocyte-like clusters, suggesting progressive differentiation from a shared progenitor state. (**B**) Along this trajectory, cells show decreasing expression of MDM2 and increasing expression of HMGA2, markers associated with DD and WD histology, respectively. (**C**) Genes dynamically regulated across pseudotime were clustered into co-expression modules, revealing distinct transcriptional programs associated with cell state transitions. One module—designated group 3—was characterized by low expression early in pseudotime and progressive upregulation toward the terminal WD-specific clusters. (**D**) Groups were analyzed for overrepresentation of transcription factor target genes.

To gain insight into the transcriptional programs driving this transition, we identified genes with significant dynamic expression across pseudotime and organized them into co-expression modules (**Fig. 4C**). One module—designated group 3—was characterized by low expression in early pseudotime and progressive upregulation toward the terminal WD-specific clusters. This pattern is consistent with a differentiation trajectory from common clusters to a mature adipocyte-like state and likely reflects genes activated during the acquisition of adipogenic identity. Notably, group 3 was enriched for known transcriptional targets of PPARG and other key regulators of adipogenesis. These findings suggest that adipogenic differentiation in WD tumors is orchestrated by a coordinated transcriptional program and that loss or failure to activate this program may underlie the dedifferentiated state in DD tumors. Given that PPARG is both necessary and sufficient for adipocyte differentiation in normal and malignant contexts, these data raise the possibility that reactivating PPARG signaling could restore differentiation capacity in DD LPS cells, offering a potential therapeutic strategy(*34*).

### PPARG2 promotes adipogenic differentiation and suppresses tumor growth in DD LPS

To explore whether PPARG activation might drive adipogenic differentiation in dedifferentiated liposarcoma (DD LPS), we first generated a PPARG gene score based on canonical PPARG transcriptional targets. This score was significantly enriched in the WD-specific adipocyte-like clusters identified in our single-nucleus RNA-seq dataset (**Fig. 5A**), consistent with the idea that PPARG activity marks a terminally differentiated state. These findings suggested that PPARG is a key regulator of differentiation in LPS and prompted us to test whether activating this pathway could reprogram DD tumor cells.

**Figure 5.**
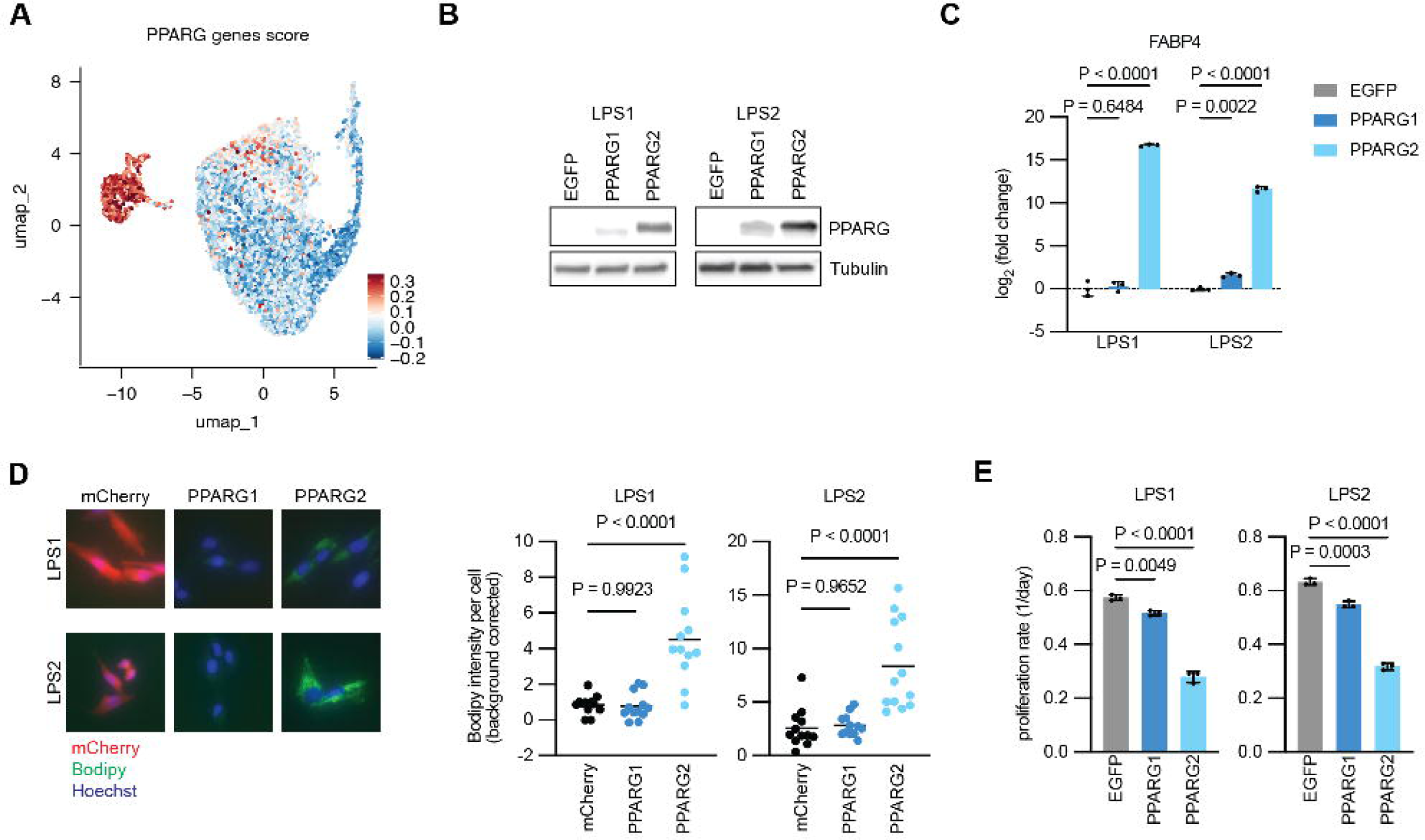
PPARG2 promotes adipogenic differentiation and suppresses proliferation in DD LPS cells. (**A**) A PPARG gene score based on known transcriptional targets is enriched in WD-specific clusters. (**B**) Western blot showing PPARG levels in DD LPS cell lines with overexpression of PPARG1 and PPARG2. (**C**) qPCR analysis shows that PPARG2 induces robust expression of FABP4, while PPARG1 has modest effects. (**D**) Bodipy staining and quantification reveal lipid droplet accumulation in PPARG2-expressing cells, consistent with adipocyte-like differentiation. (**E**) PPARG2 significantly reduces proliferation of DD LPS cell lines (LPS1 and LPS2), consistent with induction of a terminal differentiation program.

To functionally test whether PPARG activation is sufficient to induce adipogenic differentiation in DD LPS, we overexpressed its two major isoforms—PPARG1, a ligand-dependent isoform, and PPARG2, a constitutively active variant with enhanced adipogenic activity—in two DD LPS cell lines (LPS1 and LPS2) (**Fig. 5B**). Both isoforms were expressed from the same lentiviral backbone, but PPARG2 protein levels were higher, consistent with prior reports showing that PPARG2 is more stable than PPARG1 in mammalian cells(*35, 36*). Functionally, PPARG2 induced strong expression of FABP4, a canonical PPARG target gene and adipogenic marker, and promoted lipid droplet accumulation (**Fig. 5C–D**) (35). By contrast, PPARG1 had only modest effects on differentiation marker expression and lipid accumulation, suggesting that endogenous ligand availability may be insufficient to activate this isoform under baseline conditions. In addition to driving differentiation, PPARG2 expression significantly suppressed proliferation in both cell lines (**Fig. 5E**), consistent with induction of a terminal differentiation state. These findings demonstrate that PPARG2 is sufficient to both activate adipogenic gene programs and impair tumor cell growth in vitro, reinforcing its role as a central regulator of lineage fate in LPS.

To assess the therapeutic relevance of PPARG activation in vivo, we generated doxycycline-inducible LPS2 cells expressing either PPARG1 or PPARG2 and implanted them subcutaneously into immunocompromised mice (**Fig. 6A**). Tumor growth was monitored by caliper measurements, and doxycycline-containing chow was introduced once tumors reached 100–200 mm³. Induction of PPARG2 led to a significant reduction in tumor growth, while PPARG1 induction had no measurable effect (**Fig. 6B**). Histological analysis of the xenografts revealed heterogeneous tumor architecture, including the presence of lipid-rich regions reminiscent of the WD components seen in human WD/DD LPS tumors (**Fig. S5A**). Although we did not observe significant differences in the overall percentage of lipid-rich area between groups, this is likely confounded by the growth-suppressive effects of PPARG2, which limited overall tumor expansion and may have reduced the opportunity for detectable differentiation to accumulate at the tissue level (**Fig. S5B**).

**Figure 6.**
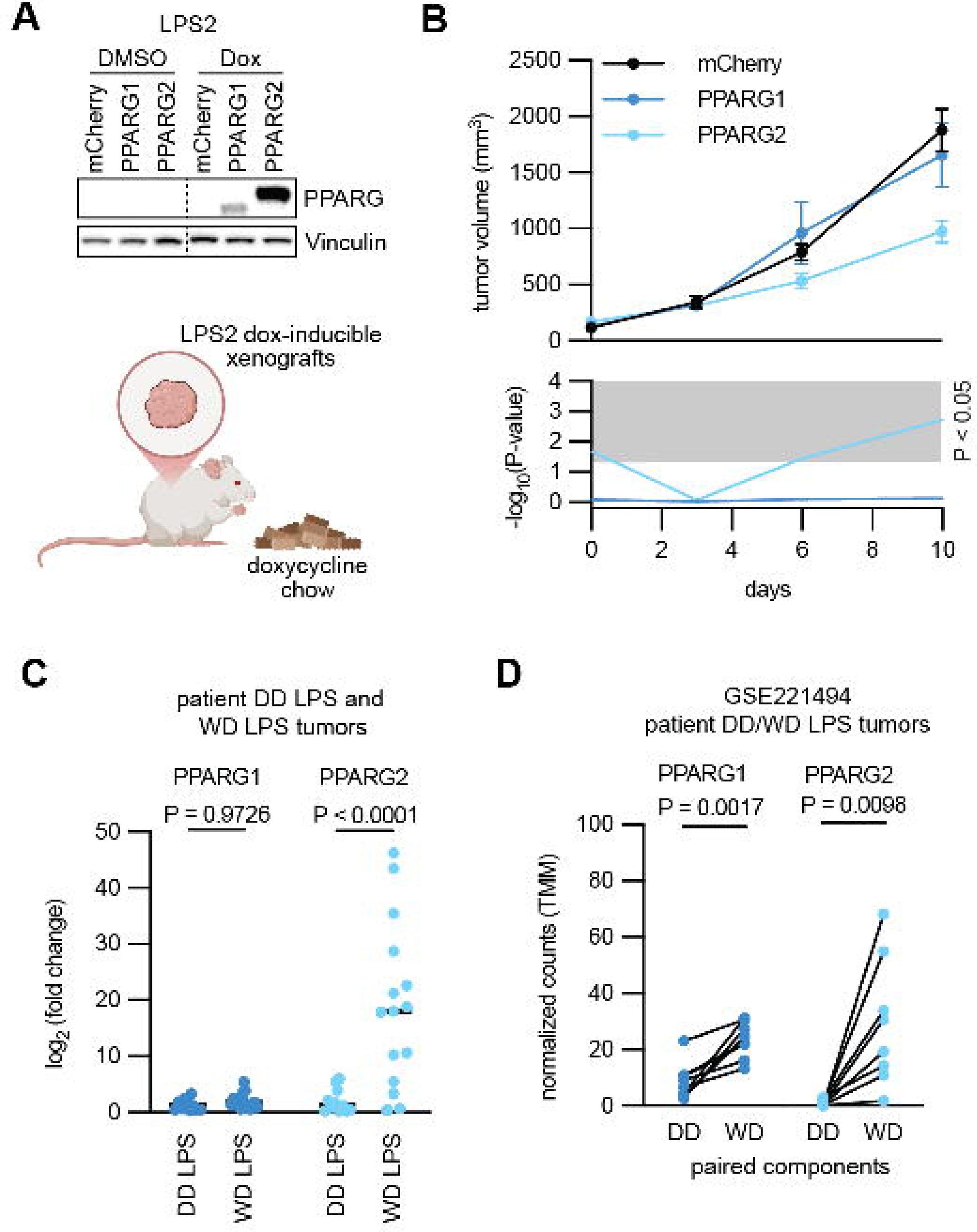
Activation of PPARG impairs tumor growth and is associated with adipogenic features in DD LPS. (**A**) Schematic of the in vivo experimental approach using doxycycline-inducible expression of PPARG1 or PPARG2 in LPS2 xenografts. (**B**) Tumor growth curves demonstrate that induction of PPARG2 significantly suppresses tumor growth, while PPARG1 has no measurable effect. (**C**) qPCR analysis of primary human tumors reveals that PPARG2, but not PPARG1, is significantly enriched in WD tumors. (**D**) Analysis of publicly available RNA-sequencing data from matched WD and DD components of WD/DD tumors confirms higher PPARG2 expression in WD regions, further supporting its role as a marker of differentiation (GSE221494)(25).

To test whether the limited activity of PPARG1 *in vivo* reflected insufficient ligand availability, we treated mice bearing parental LPS2 xenografts with rosiglitazone, a synthetic thiazolidinedione-class PPARG agonist. In adipose tissue, ligand-activated PPARG1 not only drives adipogenic gene expression but also promotes a feed-forward loop by initiating PPARG2 transcription through a conserved PPARG response element(*37*). Consistent with this mechanism, daily rosiglitazone treatment significantly slowed tumor growth (**Fig. S6A-B**), indicating that pharmacologic activation of PPARG1 can partially recapitulate the effects of PPARG2 in vivo. These findings suggest that DD LPS cells retain a latent capacity to engage the PPARG differentiation program, but fail to do so due to insufficient endogenous ligand activity. Pharmacologic activation may therefore bypass this block and re-engage differentiation pathways. While the effect size was more modest than that observed with direct PPARG2 expression, these results provide proof-of-concept that PPARG agonists can restrain tumor growth by shifting cell state, and suggest a therapeutic opportunity to restore adipogenic differentiation using clinically accessible compounds.

Finally, we examined expression of PPARG isoforms in primary human tumors. While PPARG1 levels were comparable across WD and DD samples, PPARG2 expression was significantly enriched in WD tumors and in the WD components of mixed WD/DD cases (**Fig. 6C– D, Table 2**). Together, these results support a model in which PPARG2 serves as a key driver of adipogenic differentiation in liposarcoma and suggest that restoring PPARG activity may offer a strategy to suppress proliferation and promote differentiation in DD tumors.

**Table 2.**
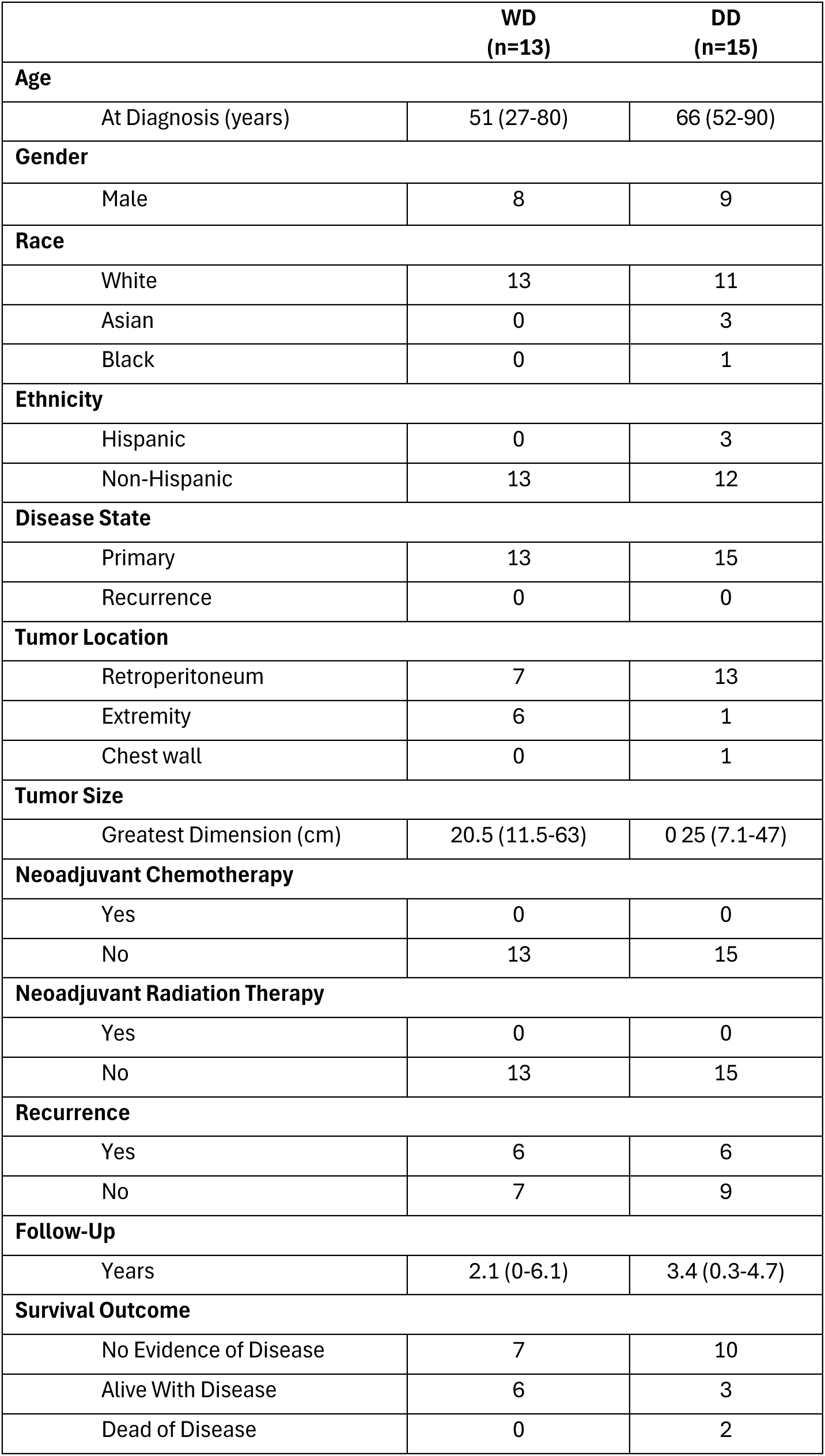
Clinicopathologic variables of WD and DD LPS tumor. Age, tumor size and follow-up are listed as median (range). Remaining variables are listed as frequency (n). Patients were diagnosed between June 2019-Aug 2024, censor date May 30 2025.

## Discussion

Well-differentiated and dedifferentiated liposarcomas (WD/DD LPS) are defined by common amplification of 12q13–15 oncogenes MDM2 and CDK4, and can co-exist in the same patient, in the same tumor, with disparate behavior along this continuum of disease, both lipoma-like indolent and high-grade rapidly fatal components. The coexistence of WD and DD lineages within a single tumor and the capacity for tumors to shift between these states over time suggests a dynamic process of differentiation and dedifferentiation. Yet, the molecular mechanisms governing these transitions remain poorly understood.

Using snRNA-seq, we defined the transcriptional landscape of spatially distinct WD and DD components from patient tumors. While most tumor cells in both compartments shared a common mesenchymal-like architecture, a rare population of adipocyte-like tumor cells was identified exclusively in WD regions. These cells expressed canonical adipogenic markers and formed a distinct transcriptional cluster, consistent with a terminal differentiation state. Pseudotime trajectory analysis revealed a unidirectional path from shared progenitor-like clusters to this adipocyte-like population, supporting a model in which the WD component may emerge from a less differentiated tumor state.

We identified PPARG as a key regulator of this transition. A PPARG gene score was enriched in the WD-specific clusters, and pseudotime-regulated genes were enriched for PPARG targets and adipogenic modules. Among the PPARG isoforms, PPARG2—a constitutively active splice variant—was sufficient to induce lipid accumulation and suppress proliferation in DD LPS cells. *In vivo*, inducible expression of PPARG2 impaired tumor growth, and pharmacologic activation of PPARG1 with rosiglitazone partially recapitulated this effect, potentially through feed-forward activation of PPARG2 expression. These findings suggest that PPARG signaling governs lineage commitment in LPS and that loss of this program contributes to the dedifferentiated state.

Importantly, our study suggests that subtype transitions in WD/DD LPS are not solely dictated by genomic divergence. Although some copy number alterations were unique to WD regions, many gene expression differences—particularly those related to differentiation—occurred independently of CNAs. These results align with recent work demonstrating that both WD and DD components arise from a common adipocyte stem cell–like progenitor and that DD tumor cells retain latent adipogenic potential(*27*). In that study, adipogenic differentiation of DD cells was actively suppressed by a TGF-β–rich microenvironment, suggesting that extrinsic cues can enforce a dedifferentiated state. Complementary work has shown that PPARG2 and its downstream targets are epigenetically silenced in DD LPS through hypermethylation of adipogenic super-enhancers(*38*). Treatment with the DNA demethylating agent 5-aza-2’-deoxycytidine and the PPARG agonist rosiglitazone restored PPARG2 expression and induced adipogenic differentiation in DD tumor cells. Together, these findings reinforce the idea that both intrinsic transcriptional programs and extrinsic environmental signals constrain differentiation in LPS and that therapeutic reactivation of PPARG2 may overcome these blocks to restore a more differentiated, less aggressive phenotype.

Together, these data support a model in which PPARG2 serves as a lineage-defining regulator in LPS, and its loss—through transcriptional suppression and/or epigenetic silencing— permits or promotes the transition to a dedifferentiated state. This model aligns with clinical behavior: WD and DD can co-exist within the same tumor or interconvert during disease progression or recurrence. Importantly, our findings suggest that subtype transitions in WD/DD LPS are not solely dictated by genomic divergence. Although some CNAs distinguish WD and DD components, many gene expression differences—particularly those related to differentiation— occur independently of copy number alterations. This further underscores the role of epigenetic and transcriptional regulation in shaping tumor cell identity.

Strategies that restore PPARG2 expression—through either demethylating agents, nuclear receptor agonists, or chromatin remodeling—may provide a means to enforce differentiation and limit aggressiveness in DD LPS. By converting a poorly differentiated, proliferative tumor into a more indolent, adipocyte-like state, it may be possible to reduce recurrence, delay progression, and improve survival. This approach mirrors successful differentiation therapies in other cancers, such as ATRA in acute promyelocytic leukemia.

More broadly, this study highlights the power of single-cell and epigenomic profiling to uncover lineage hierarchies and therapeutic vulnerabilities in mesenchymal tumors. WD/DD LPS has long presented a clinical paradox, with tumors able to recur as more or less differentiated subtypes. Our findings provide a mechanistic framework to explain these transitions and nominate PPARG2 as a therapeutic entry point for differentiation-based strategies. Understanding how PPARG activity is regulated—and how it can be restored—will be essential for translating this approach into clinical benefit.

## Materials and Methods

### Experimental Design

The objective of this study was to define cellular heterogeneity, differentiation states, and subtype transitions in well-differentiated and dedifferentiated liposarcoma (WD/DD LPS) using single-nucleus RNA sequencing (snRNA-seq). We aimed to uncover lineage relationships between WD and DD tumor components, identify molecular features of dedifferentiation, and reveal potential therapeutic vulnerabilities. To achieve this, we collected freshly resected tumor specimens from patients with histologically confirmed WD/DD LPS. Tumors with both WD and DD regions were macrodissected and processed independently to preserve distinct transcriptional programs. Nuclei were isolated and profiled using the 10x Genomics Chromium platform. Sequencing data were analyzed using a standardized computational pipeline for quality control, dimensionality reduction, clustering, and differential gene expression.

### Patients

This study was conducted in accordance with the principles expressed in the Declaration of Helsinki. Informed consent was provided by all patients or guardians granting access to tumor tissue, serum and medical records. Databases including patient identifiers were maintained according to our local institutional guidelines. The research was conducted under protocol #10-001857, approved by the UCLA Institutional Review Board (IRB).

Patients with primary, untreated WD/DD LPS undergoing initial surgical resection were selected for inclusion. Preoperative imaging was used to assist with patient selection. Patients with history of systemic chemotherapy, radiation or prior resection were excluded.

Clinicopathologic data was collected and stored in an encrypted database and maintained by the study authors (KK, BK).

### Tumor harvest, storage and validation

Fresh tissue was harvested at the time of surgery from both WD and DD components and stored immediately in liquid nitrogen. Additional tissue from both components was placed in formalin for permanent fixation and then embedded in paraffin. Tumor diagnosis and subtype was validated by expert histological review and DNA FISH (Empire Genomics, SKU MDM2-CHR12-20-GROR) was used to confirm the presence of *MDM2* amplification (authors SD, MN). Methods have been previously described(*39*). Tumors without *MDM2* amplification were excluded.

### Immunohistochemistry

Formalin-fixed sections embedded in paraffin were cut at 4 μm thickness. Samples were submerged in xylene for paraffin removal and then rehydrated using graded ethanol washes. Sections were counterstained with hematoxylin and eosin. Brightfield slides were digitally scanned on a ScanScope AT2 (Leica Biosystems, Vista, CA, USA) and analyzed using QuPath version 0.2.3.

### Nuclei isolation and cDNA library generation

After tumor diagnosis and subtype were confirmed, samples were thawed in RNA*Later*^TM^-ICE (ThermoFisher) overnight at −20°C. Nuclei were then isolated using the Chromium Nuclei Isolation with RNase Inhibitor Kit (10X Genomics). For DD components, 50mg of tissue was used and for WD components, 250mg of tissue was used from each sample. Nuclei concentration and cell viability was determined using a Countess II FL Automated Cell Counter (ThermoFisher). Cell viability <5% was used as a threshold to ensure high quality nuclei isolation. Samples were also assessed by brightfield microscopy for the presence of significant debris. Single nuclei suspensions generated from each sample were then used to construct 3’GEX cDNA libraries (10x Genomics).

### Single nuclei sequencing and processing

Single nuclei suspensions generated from each sample were used to construct 3’GEX cDNA libraries (10x Genomics) followed by next-generation sequencing via NovaSeq 6000 (Illumina). Demultiplexed sequencing results were aligned to the reference genome and ambient RNA was removed using CellBender. After filtering, 27,333 nuclei were included in the computational analysis pipeline. The R package Seurat was applied to integrate samples, cluster cells, and identify differentially expressed genes. LPS clusters were analyzed by pseudo-time analysis (Monocle3) to establish trajectory inferences between WD and DD subtypes.

### Cell lines

LPS1 and LPS2 are derived from DD LPS patient-derived xenografts, which have been previously validated(*40*). Both lines were cultured in DMEM with 10% FBS and penicillin/streptomycin, and maintained in a 37°C humidified, normoxic chamber supplemented with 5% CO_2_. Cells were monitored regularly for the presence of mycoplasma.

### Plasmids

TFORF3549 (pLX317-EGFP) was a gift from Feng Zhang (Addgene plasmid # 145025 ; http://n2t.net/addgene:145025 ; RRID:Addgene_145025). TFORF3550 (pLX317-mCherry) was a gift from Feng Zhang (Addgene plasmid # 145026 ; http://n2t.net/addgene:145026 ; RRID:Addgene_145026). TFORF3138 (pLX317-PPARG1) was a gift from Feng Zhang (Addgene plasmid # 144614 ; http://n2t.net/addgene:144614 ; RRID:Addgene_144614). TFORF3139 (pLX317-PPARG2) was a gift from Feng Zhang (Addgene plasmid # 144615 ; http://n2t.net/addgene:144615 ; RRID:Addgene_144615). pRSV-Rev was a gift from Didier Trono (Addgene plasmid #12253; http://n2t.net/addgene:12253; RRID:Addgene_12253). pMDLg/pRRE was a gift from Didier Trono (Addgene plasmid #12251; http://n2t.net/addgene:12251; RRID:Addgene_12251). pCMV-VSV-G was a gift from Bob Weinberg (Addgene plasmid #8454; http://n2t.net/addgene:8454; RRID:Addgene_8454).

Plasmids for inducible expression of mCherry, PPARG1, and PPARG2 were generated using standard molecular cloning techniques (pCW57.1-mCherry, pCW57.1-PPARG1, and pCW57.1-PPARG2).

### Lentivirus Production and Transduction

Lentivirus was produced in HEK293T cells infected with pLX317 transfer plasmid, pRSV-Rev, pMDLg/pRRE, and pCMV-VSV-G. At 2 days post-transfection, supernatant was collected and filtered at 0.45 µm, distributed into 1 mL aliquots, then froze at −80C for future use.

For transduction, aliquots of virus were thawed and polybrene was added to 4 µg/mL. The virus/polybrene mixture was added to PBS-rinsed cells and incubated for 16 hours. At that point the virus/polybrene was replaced with fresh media. After 24 hours, selection was started by treating cells with 2 µg/mL puromycin.

### RT-qPCR

For cell lines, total RNA was harvested from approximately 300,000 cells. Cells were washed with ice cold PBS then washed with RNAlater (Invitrogen). After aspirating the RNAlater, cells were frozen at −80C for processing at a later point. Cells were thawed and total RNA was collected using a Direct-zol RNA Miniprep kit (Zymo).

For patient tumors, approximately 10 mg was incubated in RNAlater-ICE (Invitrogen) overnight at −80 °C. The tumor was removed from the RNAlater-ICE, and homogenized in 1 mL trizol. Total RNA was isolated from the homogenate using RNeasy lipid tissue kit (Qiagen).

For both cell lines and patient tumors 1 µg RNA was reverse transcribed using iScriptTM Reverse Transcription Supermix (BioRad). qPCR reactions were up using Power SYBR Green PCR Master Mix (Applied Biosystems). For *PPARG1* and *PPARG2* measurements, unique forward and a common reverse primers were used – PPARG1-fwd: 5’-GCCATTTTCTCAAACGAGAGTCAGCC-3’; PPARG2-fwd: 5’-TGACCCAGAAAGCGATTCCTTCA-3’; PPARG1/2-rev: 5’-ACGGAGAGATCCACGGAGCTGA-3’. For FABP4, the following primers were used – FABP4-fwd: 5’-ACGAGAGGATGATAAACTGGTGG-3’; FABP4-rev: 5’-GCGAACTTCAGTCCAGGTCAAC-3’.

### Lipid content quantification

Live cells were stained with 1 µM Bodipy 493/503 (Invitrogen) and 0.2 µg/mL Hoechst-33342 (Invitrogen). Cells were incubated in media containing dyes at 37°C for 10 min, then washed well three times with PBS. Cells were imaged using an Evos Cell Imaging System.

QuPath was used to analyze images for fluorescent intensity. In short, 5 cells from each sample and 3 samples from each condition were analyzed for a total of 15 cells per condition. Mean Bodipy fluorescent intensity was determined from each region of interest.

### Proliferation assays

Cells were seeded at a density of 20,000 cells/well in 6-well dishes. After 24 hours cells were counted and used as day 0 measurements. At day 3 cells were counted from separate wells and the data was fitted to an exponential growth model. The fitted growth rates were presented as the proliferation rates with units of divisions/day.

### Mouse xenografts

LPS2 cells were used to establish tumor xenografts. A total of 1,000,000 cells in a 1:1 mixture of Matrigel and PBS were injected into the one side of 7-week-old NSG mice. Tumors were monitored for growth by calipering. Tumor volumes were calculated by the equation (length x width^2^)/2. When tumors reach 100-200 mm^3^, treatment was initiated. Rosiglitazone was administered via oral gavage daily at a concentration of 20 mg/kg. Mice were weighed daily and exhibited no significant weight loss due to rosiglitazone treatment. Mice were euthanized once tumors reached 2000 mm^3^.

LPS2 cells expressing doxycycline-inducible mCherry, PPARG1, or PPARG2 were used to establish tumor xenografts. A total of 1,000,000 cells in a 1:1 mixture of Matrigel and PBS were injected into the one side of 7-week-old NSG mice. Once tumors reached 100-200 mm^3^ all mice were fed chow containing 625 mg/kg doxycycline hyclate (Envigo). Mice were euthanized once tumors reached 2000 mm^3^.

### Statistical Analysis

Statistical significance was established using a *P* value threshold < 0.05 with 95% confidence intervals. Continuous, normally distributed data were evaluated using the two-tailed T-tests for pairwise comparisons and ANOVA for comparisons involving multiple groups. The normality of the data was confirmed using quantile-quantile (Q-Q) plots. The Bonferroni test was employed for post-hoc analysis to identify specific group differences. Data lacking normal distribution was assessed using the Mann-Whitney U test. All experiments were carried out in duplicate or triplicate to ensure reliability. Statistical analyses were performed using Graphpad Prism software, version 9.3.1, on a MacOS platform. Unless otherwise indicated, data are reported as mean ± standard deviation (SD).

## Acknowledgments

We thank the members of the Christofk and Kadera labs for their discussion and constructive feedback.

## Funding

UCLA Jonsson Comprehensive Cancer Center (JCCC) seed grant (BEK and KDK)

UCLA Department of Surgery grant (BEK and KDK)

NIH R01 CA215185 and R01 CA215185 and R01 AR070245 (HRC)

American Cancer Society 133839-PF-19-203-01-CCG (BRW)

## Author contributions

Conceptualization: BRW, KDK, BK, HRC Methodology: BRW, KDK, BK, HRC Investigation: BRW, KDK, FD, CD, CF Supervision: BK, HRC

Writing—original draft: BRW, KDK Writing—review & editing: BRW, KDK, BK, HRC

## Competing interests

Authors declare that they have no competing interests.

## Data and materials availability

All data, code, and materials used in the analyses must be available in some form to any researcher for purposes of reproducing or extending the analyses. Include a note explaining any restrictions on materials, such as materials transfer agreements (MTAs). Include accession numbers to any data relevant to the paper and deposited in a public database; include a brief description of the dataset or model with the number. The DMA statement should include the following: “All data are available in the main text or the supplementary materials.”

---

**Fig. S1.**
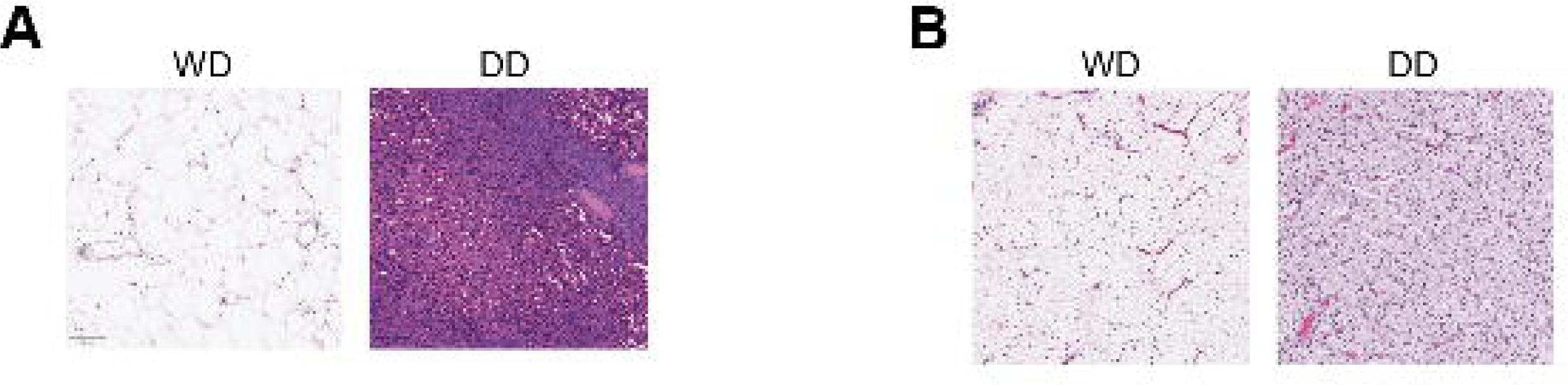
Histologic characterization of WD and DD tumor components. (**A**-**B**) H&E staining of two WD/DD LPS tumors used in snRNAseq shows consistent histological differences between WD and DD regions, with WD components displaying abundant mature adipocytes and lipoblasts, and DD regions exhibiting dense cellularity and spindle morphology.

**Fig. S2.**
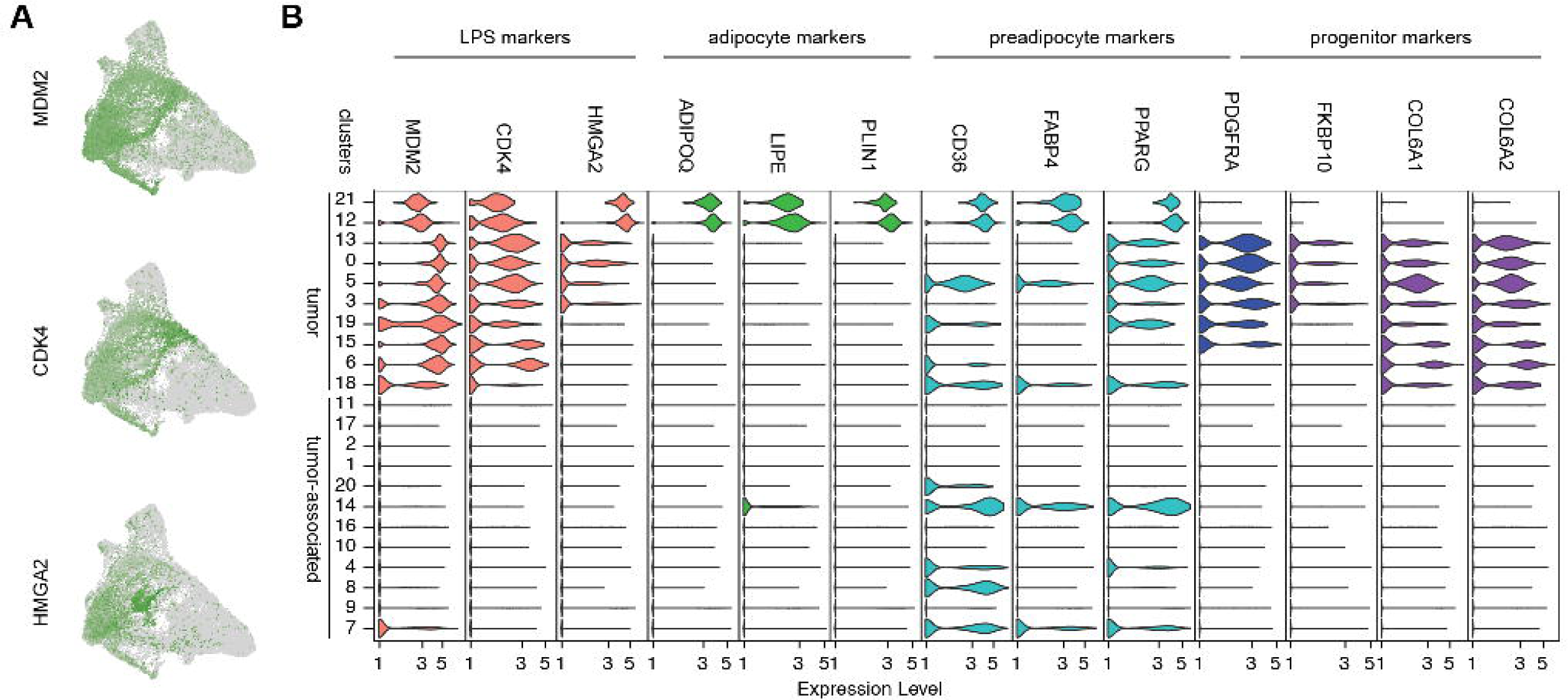
Identification of tumor clusters based on expression of LPS oncogenes and copy number alterations. (**A**) UMAP feature plots for *MDM2*, *CDK4*, and *HMGA2* across all nuclei. (**B**) Violin plots further confirm selective expression of these oncogenes as well as adipocyte, preadipocyte, and adipocyte progenitor markers.

**Fig. S3.**
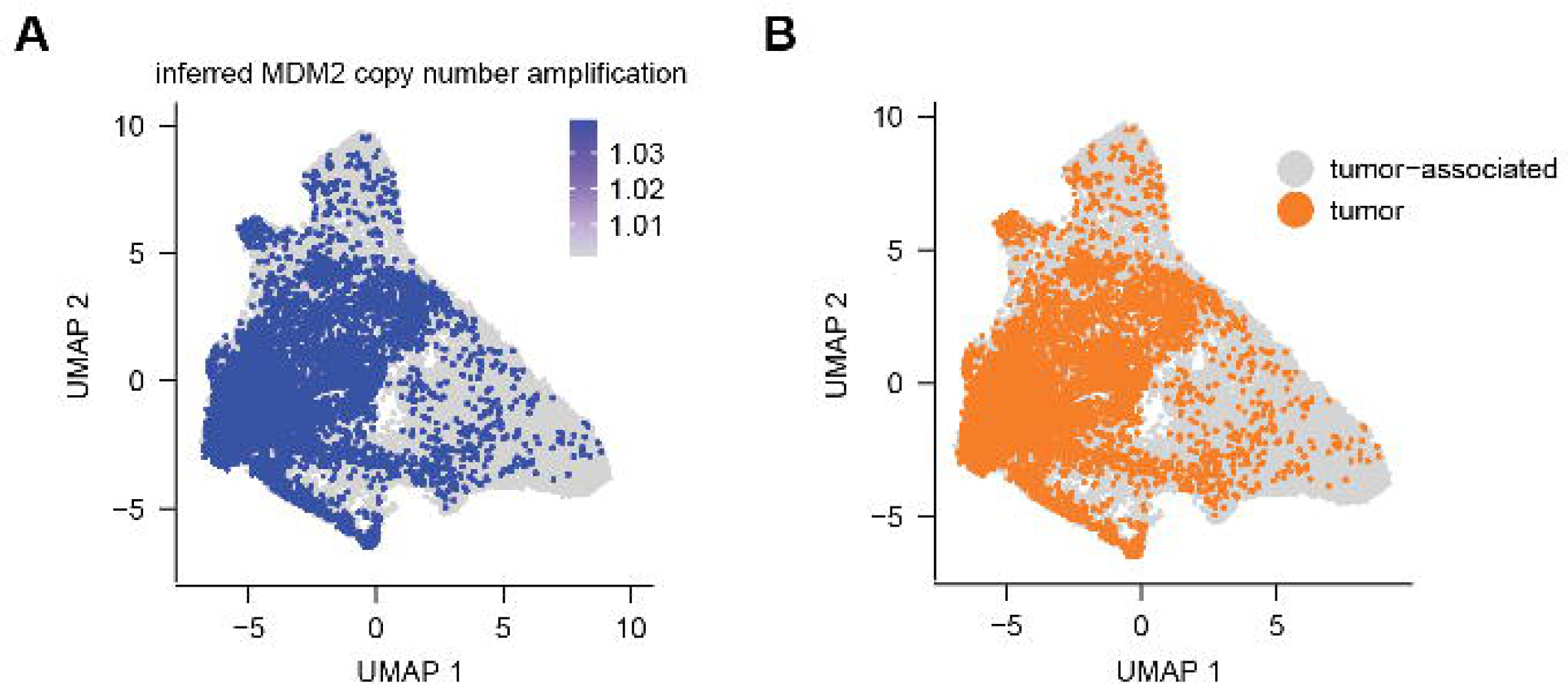
Inferred copy number gains in MDM2 are used to identify tumor cells. UMAP profiles across all clusters highlight (**A**) amplification of MDM2 and (**B**) annotation of tumor cell populations.

**Fig S4.**
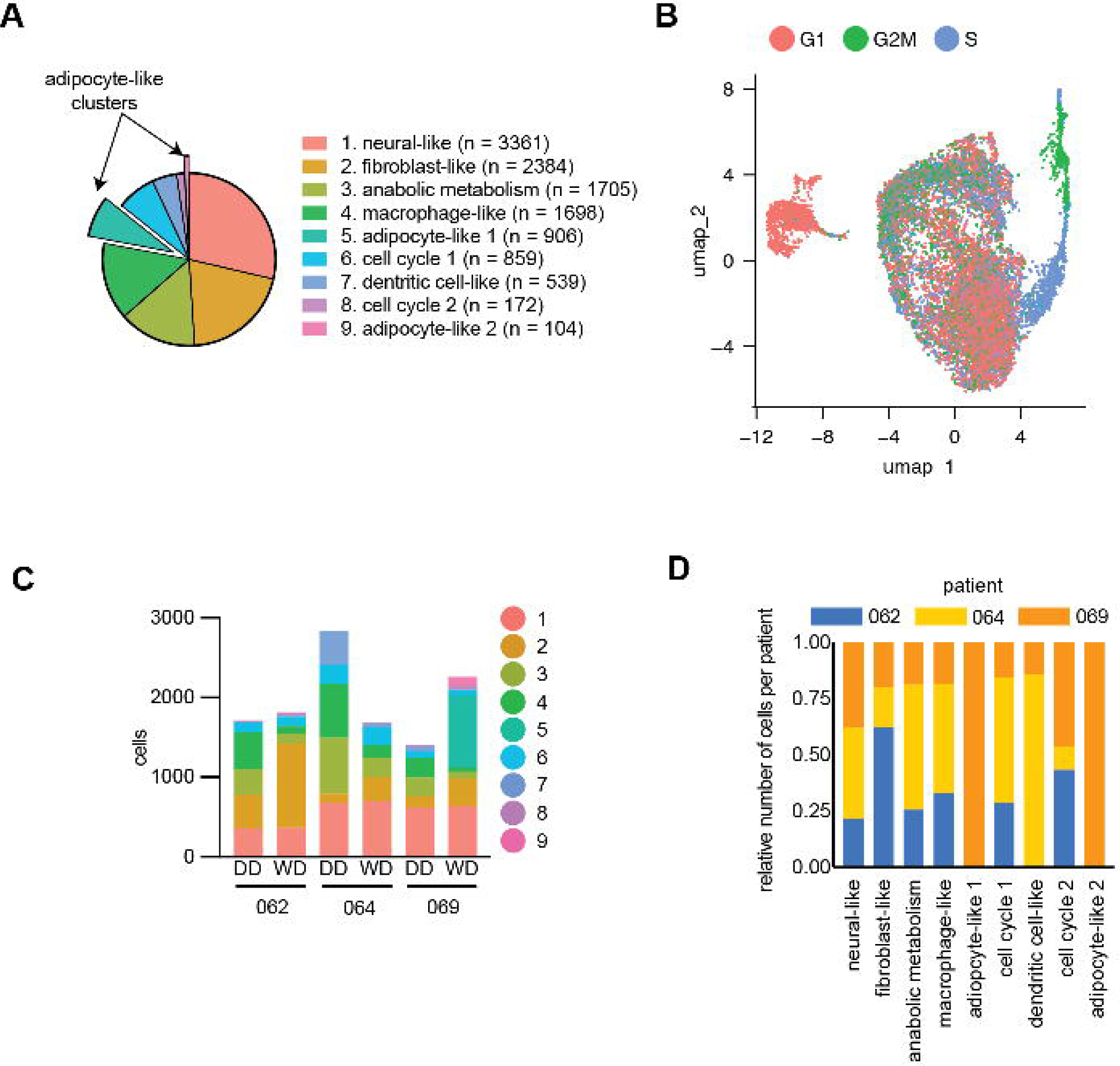
Functional annotation of re-clustered tumor cell states. (**A**) Pie chart shows relative number of nuclei across the nine malignant clusters, including corresponding to proliferative, metabolic, mesenchymal-like, immune-like, and adipocyte-like states. (**B**) UMAP plots colored by predicted cell cycle state of each nuclei. (**C**) Number of nuclei contributing to each cluster, split by patient and component. (**D**) The relative number of nuclei per patient for each cluster.

**Fig S5.**
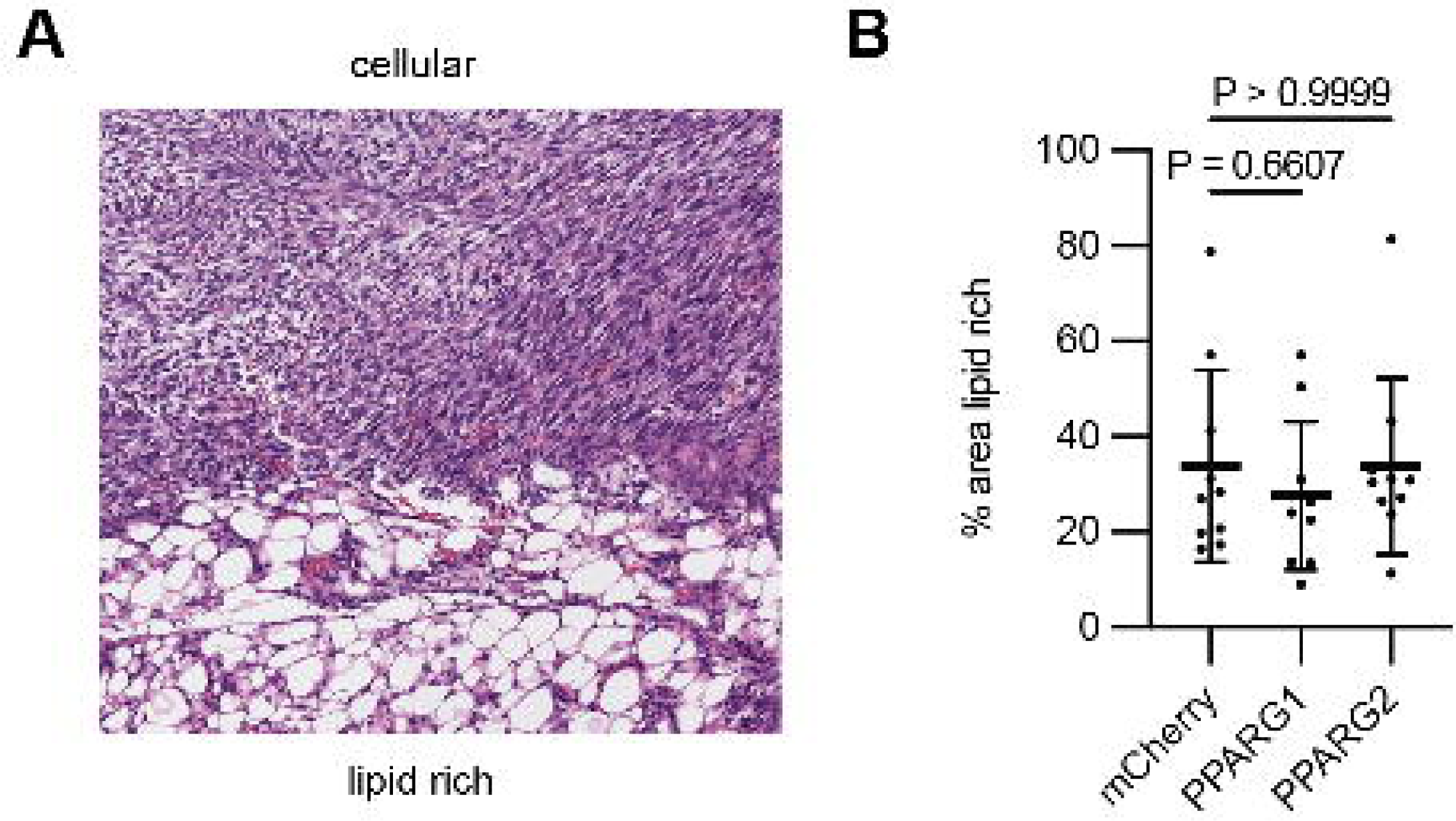
Histological analysis of LPS2 xenografts with induction of PPARG isoforms. (**A**) Representative H&E staining of LPS2 xenograft tumors following doxycycline-induced expression of PPARG1 or PPARG2 reveal focal lipid-rich regions. (**B**) Quantification of lipid-positive area shows no differences, likely due to limited tumor size following PPARG-mediated growth suppression.

**Fig S6.**
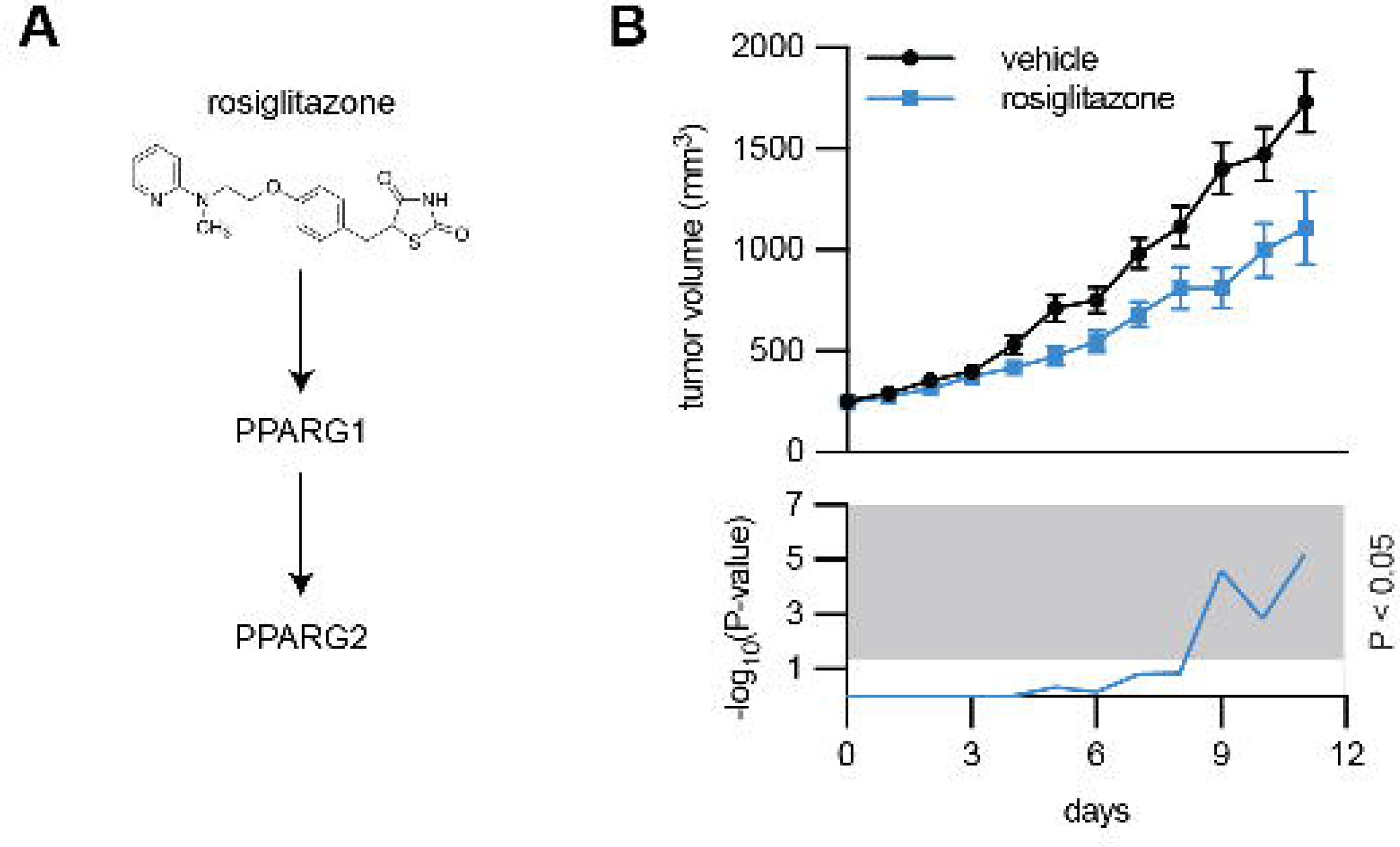
PPARG activation decreases DD LPS tumor growth in vivo. (**A**) Rosiglitazone activates PPARG1 activity, which drives expression of PPARG2. (**B**) Rosiglitazone treatment of mice bearing parental LPS2 xenografts reduces tumor growth, supporting the idea that spharmacologic activation of PPARG1 can partially recapitulate the effects of PPARG2 *in vivo*.

## Notes

### Competing Interest Statement

The authors have declared no competing interest.

